# RAFTS3: Rapid Alignment-Free Tool for Sequence Similarity Search

**DOI:** 10.1101/055269

**Authors:** Ricardo Assunção Vialle, Fábio de Oliveira Pedrosa, Vinicius Almir Weiss, Dieval Guizelini, Juliana Helena Tibaes, Jeroniza Nunes Marchaukoski, Emanuel Maltempi de Souza, Roberto Tadeu Raittz

## Abstract

**Background:** Similarity search of a given protein sequence against a database is an essential task in genome analysis. Sequence alignment is the most used method to perform such analysis. Although this approach is efficient, the time required to perform searches against large databases is always a challenge. Alignment-free techniques offer alternatives to comparing sequences without the need of alignment.

**Results:** Here We developed RAFTS3, a fast protein similarity search tool that utilizes a filter step for candidate selection based on shared k-mers and a comparison measure using a binary matrix of co-occurrence of amino acid residues. RAFTS3performed searches many times faster than those with BLASTp against large protein databases, such as NR, Pfam or UniRef, with a small loss of sensitivity depending on the similarity degree of the sequences.

**Conclusions:** RAFTS3 is a new alternative for fast comparison of proteinsequences genome annotation and biological data mining. The source code and the standalone files for Windows and Linux platform are available at: https://sourceforge.net/projects/rafts3/

## BACKGROUND

Biological data mining deals with the discovery of patterns, trends, answers, or other meaningful information that is hidden in the data. Sequence comparison is the main component in the retrieval system from genomic databases. An efficient sequence comparison algorithm is critical for searching biological databases. Usually, bioinformatics workflows use algorithms based on sequence alignment such as BLAST [1] to search for similarity of DNA/RNA or protein sequences against large sequence databases. Comparisons involving large databases such as NCBI NR [2],however, are computationally costly and demand long running times. The development of new computationally faster algorithmsmay providesignificant improvement in biologicalpattern search. A class of techniques that can speed up sequence comparison is the alignment-free approach [3].

Algorithms based on sequence alignment are efficient in detecting similarities between protein sequences. These approaches have been improved since the first methods. Originally alignment techniques used dynamic programming to produce an optimized alignment between the sequences. Although efficient implementations have been developed, the computational load to compare large amounts of sequences makes these algorithms very slow and demanding [3, 4]. To compensate for the high computational cost of full alignments, heuristic approaches were proposed. In general, these methods use subsequences of pre-determined length "k" (k-mers). The subject database is searched to find sequences that have common k-mers related to the query sequence. The k-mers are then extended using scores schemes to maximize the aligned regions. However, although heuristic methods are somewhat efficient to perform searches in large databases, they also have their limitations, such as loss of sensitivity and parameter thresholds [4].

The alignment-free methods offer a way to obtain a similarity measure between sequences without the need to perform alignments. These methods are also based on the assumption that two similar sequences share a certain portion of k-mers. Given a query sequence, the alignment-free methods generally work by selecting subject sequences with k-mers that are present in both query and subject sequences. The procedure then applies a statistical method to establish a similarity ranking for these sequences [5].

Generally, alignment-free techniques are divided in two classes: a) methods based on words (sequences) with fixed sizes, followed by the use of statistical analysis including procedures based on defined metrics such as Euclidean distance and entropy of frequency distributions; and b) methods where words of fixed sizes are not required for statistical analysis, using data compression and/or Kolmogorov complexity scale independent representations by iterated maps. Reviews of these techniques are available at [3, 5, 6].

Several alignment-free techniques have been proposed with different degrees of success. The very first proposal of an alignment-free method for biological sequence comparison showed to be superior to alignment based algorithms in some aspects such as the ability to compare low similarity sequences [7].Since then, it has been applied in phylogenetic reconstruction [8–11], identification of homologous proteins [4], genome annotation [12], classification of metagenomic sequences [13], and identification of regulatory sequences [14]. Also, it has been shown as an efficient technique for sequence filtering [15].

Alignment-free approaches have been used to replace alignment based approaches for searching and comparing sequences against large databases showing significant increase in speed. PAUDA [16] is an alternative to BLASTx for searching sequencing reads against protein databases in metagenomics. PVC (Periodicity Count Value) is a method for finding homologous nucleotide sequences as alternative for BLASTn [17]. USEARCH [18] is an alternative to BLASTp that applies a k-mer approach to perform searches of protein sequences against a protein database.

In this paper we propose a fast and efficient alignment-free method named RAFTS3. The method is based on amino acid co-occurrence matrices and on a new heuristic approach for filtering sequences. The results show that RAFTS3 is much faster than BLASTp with negligible loss of sensitivity when applied against large databases in all tests performed and can be successfully used in several biological data-mining tasks.

## IMPLEMENTATION

Since RAFTS3 deals with protein sequence comparison against protein databases, the first step to be considered is to set up the protein database into a specific RAFTS3 format. The formatting consists of two steps to be applied to each protein sequence within a FASTA file: a) the sequences must be indexed by a hash function and b) a binary amino acid co-occurrence matrix (BCOM) has to be assigned to each sequence to represent its contents.

When a formatted database is available, query searches can be performed. This process is also divided in two distinct steps: (1) the filtering of candidates, that selects sequences whose indexed k-mers are shared with the query sequence, and (2) the comparison of these candidates, that is done by means of the BCOM.

### Database formatting process

The formatting process takes a FASTA database as input and creates a file comprising a hash table and the BCOM matrices for all sequences in the database. Aiming to improve access to the sequences, RAFTS3 also creates an index to allow direct access to each sequence in the FASTA file (Figure 1.A).

**Figure 1.**
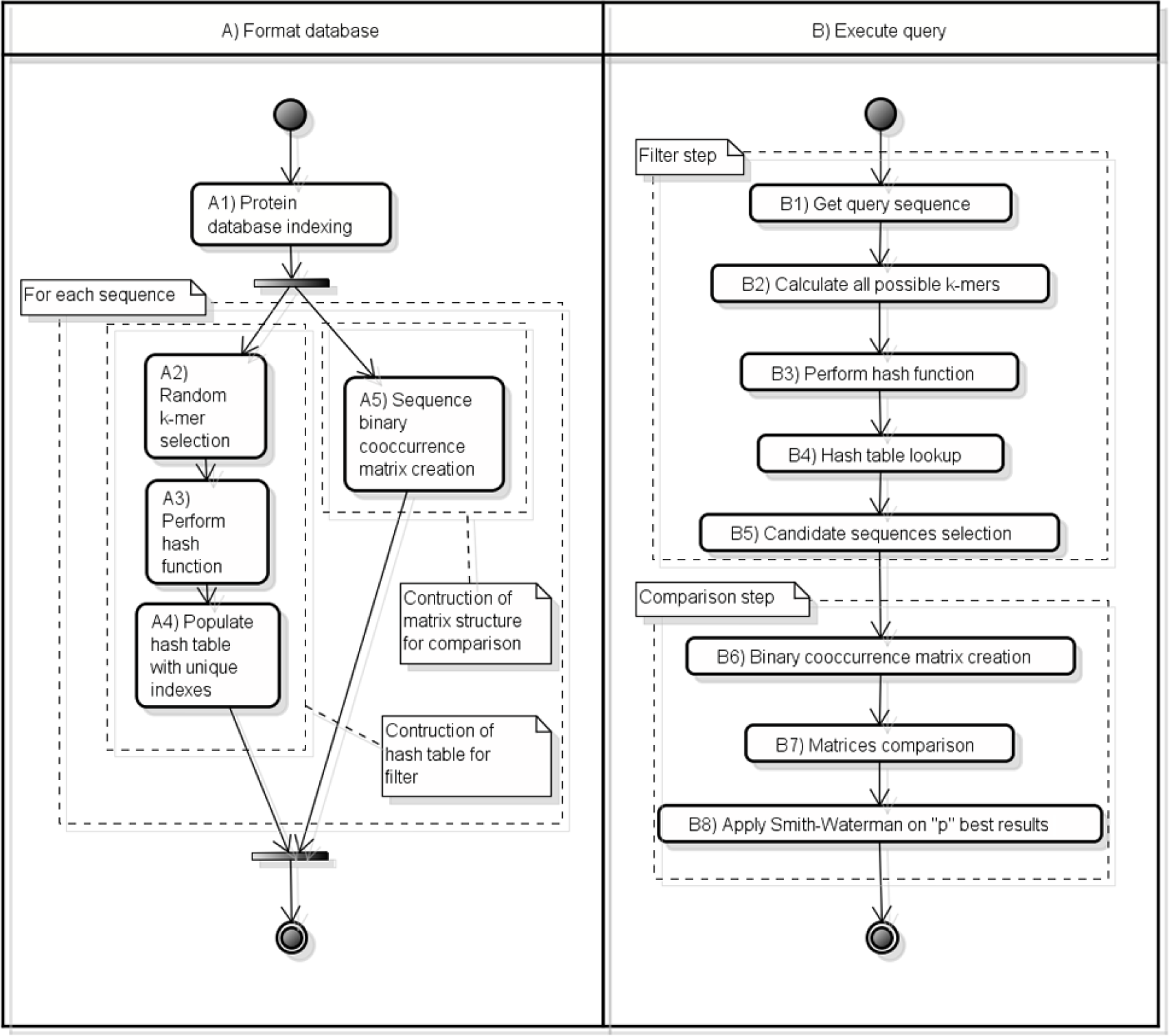
**RAFTS3 activity diagram.** RAFTS3 format database and query search overview. A) Shows the database formatting processes, which involve construction of two structures used in query sequence search, a hash table and a set of binary co-occurrence matrices. B) Shows the process for searching and comparison of a query sequence, with filtering and comparison steps separated.

For each sequence in the database a set of k-mers is randomly selected and submitted to a hash function. The indexes are then stored into a hash table for fast selection of candidate for comparison. These indexes will permit further retrieval of any sequence in the database sharing a given k-mer. As default, 10 k-mers with lengths of 6 amino acid residues are selected per sequence.

The formatting process also involves a BCOM assignment to each sequence. The BCOM was designed to represent the sequences using few bytes of memory. Both the hash table and the BCOM matrices are stored in a common structure that is loaded in RAM with the application aiming to minimize disk access when comparing sequences. The hash function and the BCOM structure will be detailed further.

### Query sequence search

Searching is the goal step in RAFTS3. Its purpose is to retrieve similar sequences to a sequence of interest from a database. Also, it is desirable that the recovered sequences are ranked by their similarity with the query sequence. Searching involves two main steps: filtering and comparison.

In the filtering process, the search scope is reduced by selecting, through a hash table, only sequences containing common k-mers related to the query sequence. To perform a search based on a sequence of a given length *n*, hash indexes forall possible k-mers with length *k* are calculated by taking a sliding window that runs through the sequence from position 1 to *n*−*k*+ 1. The indexes generated for each k-mer are used to select the candidate sequences by consulting the hash table (Figure1.B).

The comparison is performed with the candidate sequences based on their BCOM. The details of the comparison method will be discussed later (see Binary cooccurrence matrix (BCOM)). Alignments of the best results can also be done to confirm the results or to assign them to a well-established metric. The number of alignments can be customized by parameters; by default, a Smith-Waterman alignment [19] is performed only with the best stated result. As a measure of alignment quality, besides the alignment score, we calculate a relative score (E (1) [20]:

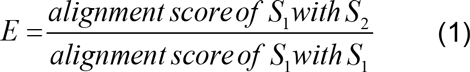
where *S*_1_ and *S*_2_ are protein sequences. E-values are also computed using Karlin Altschul statistics [21].

### Hash function for candidate sequence selection

The hash function of RAFTS3 is an essential step in the filtering process and it is applied to both database and query. The recursive indexing technique (INREC) [22] was used to assign a real number to a protein k-mer. INREC is a technique of dimensionality reduction and pattern recognition that uses arecursive process of a mathematical function to encapsulate, in a single number, the information that describes a pattern. Thereby, the indexes generated by similar sequences are equal or close to each other. The numbers generated by the INREC function are transformed in hash indexes *H* through the expression (2).

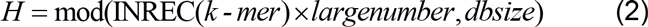

Where *largenumber* is a value to express the decimal fraction of the INREC index as an integer number, and *dbsize* defines size and spreading of the hash table. By using the hash table, sequences sharing the same INREC indexes are rapidly selected as candidate for comparison.

To apply the INREC algorithm, amino acid residues need to be converted to a quaternary numeral system triplet by a two-way conversion table (Table 1). The numbers are arbitrary, but the codes are assigned in correspondence to possible codons. The numerals1, 2, 3 and 4 represent the nucleotide residues A, C, G and T/U, respectively.

**Table 1.** Amino acid numeric conversion

| Amino acid | Code |
| --- | --- |
| A | 3 2 4 |
| R | 1 3 1 |
| N | 1 1 2 |
| D | 3 1 2 |
| C | 4 3 2 |
| Q | 2 1 1 |
| E | 3 1 1 |
| G | 3 3 4 |
| H | 2 1 4 |
| I | 1 4 1 |
| L | 2 4 2 |
| K | 1 1 1 |
| M | 1 4 3 |
| F | 4 4 2 |
| P | 2 2 4 |
| S | 4 2 1 |
| T | 1 2 4 |
| W | 4 3 3 |
| Y | 4 1 2 |
| V | 3 4 2 |

Two-way conversion table of amino acid residues to quaternary numeral system triplets.

Thus, given a sequence of integers *D*={*d*_1_, *d*_2_,…*d_m_*} representing a sequence of length *m*, where *d*_1_ ∈(1,2,3,4}. The INREC index *I* is generated from the recursion of the function f:

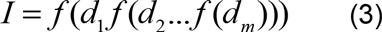
where 
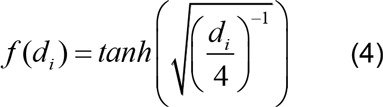

The amino acid sequence *MAF* can be used to illustrate how the indexing works. By using the conversion table (Table 1) the amino acids are represented as *M*= {1,4,3}, *A*= {3,2,4} and *F*= {4,4,2}. Thus, the sequence *MAF* can be represented as *D* ={1,4,3,3,2,4,4,4,2}. Applying the *f* function recursively (4 and 3), from the last element to the first: 
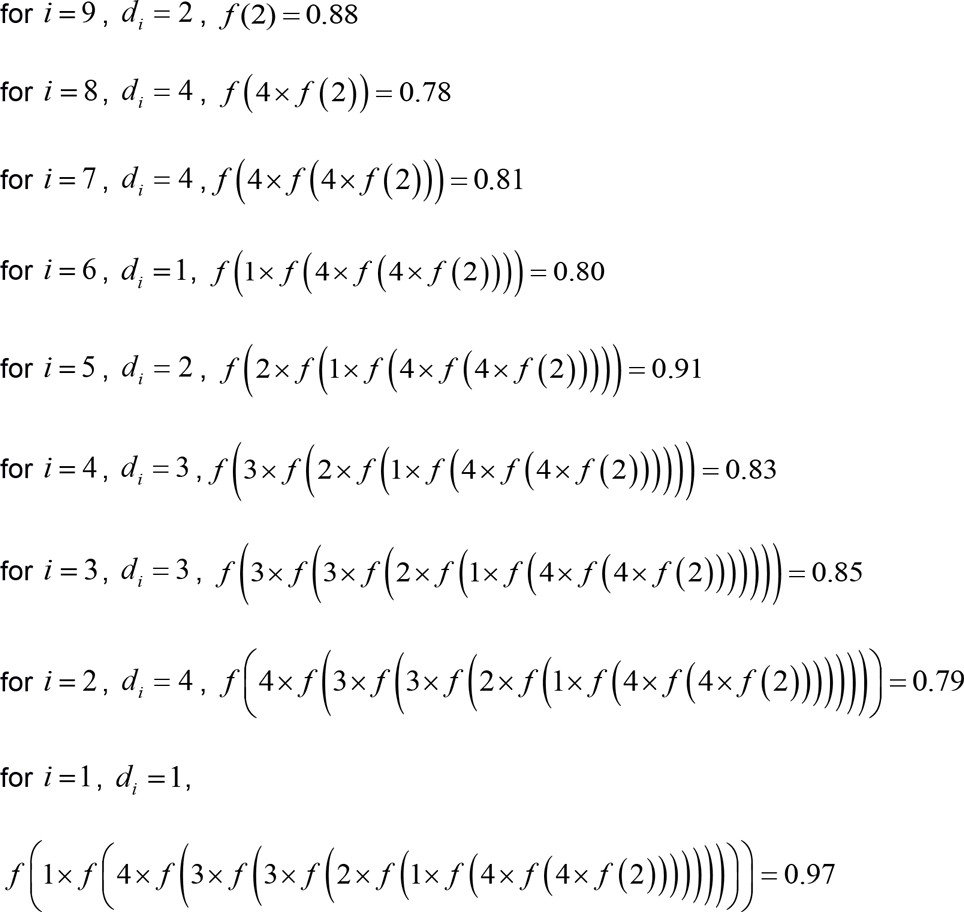

Therefore, for the sequence MAF, the INREC index *I* is 0.97.

### Binary co-occurrence matrix (BCOM)

The binary co-occurrence matrix BCOM is a bi-dimensional fingerprint of an amino acid sequence. It not only represents an amino acid sequence but is a pattern for comparison with other sequences.

A BCOM is a binary matrix where each cell position (*x, y*) represents the occurrence of an amino acid pair *XY* in a sequence*S*. If the value within the cell is set to null, the pair does not occur in *S* (Figure 2). Thus for each sequencea 20×20 binary matrix is generated representing the occurrence of all possible amino acid pairs within it. Thereby, any sequence can be represented by a matrix with 400 bits or 50 bytes. The small data volume and the uniform structure of the BCOM allows databases with millions of sequences to be represented and stored in RAM. The entire NR database can be handled in a common laptop.

To compare two matrices, let *A* and *B* be BCOMs corresponding to sequences *S*_1_ and *S*_2_ respectively. The binary sum between the matrices *A* and *B* represents the occurrence of common amino acid residue pairs and reflects the sequences similarity. Similarly, the binary operation *xor* is performed to calculate the degree of dissimilarity as a support for the comparison. Thus, the measure of difference e between *A* and *B* is given by the equation (5).

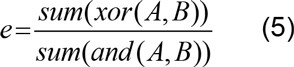

Each candidate sequence selected in the filter step is related to a dissimilarity measure given by *e*. Finally, correlation coefficients *r* (6) between the matrices are also calculated for BCOMs of sequences with highest similarity based on *e* and are used for reordering the results. Correlation coefficients are usually used to compare image differences; here the same was done with the BCOMs as an estimate of sequence identity. For instance, the sequence of the major facilitator superfamily protein of *Serratia* sp. *AS12* (gi 333925879) shares about 80% identity with the arabinose efflux permease family protein of Rahnella aquatilis (gi383191252) and the correlation coefficient is 73%; in contrast the amino acid transporter of *Aspergillus oryzae* shares about 20% of identity with the former while the correlation coefficient is 28%.

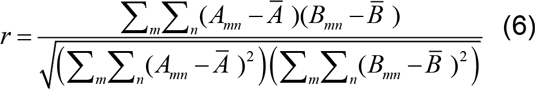

Where, Ā=*Mean*(*A*) and 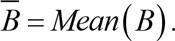

Due the computational cost, the number of sequences compared with the correlation equation (6) was limited to 50.

### Implementation and datasets

RAFTS3 was written in MATLAB using its built-in functions, the Bioinformatics Toolbox [23] and an in-house library. Three protein databases were used, the NCBI NR with 19,689,576 sequences, PFAM [24] with 15,929,002 sequences and the UniRef50 [25] with 6,784,251 sequences. The performance and sensitivity of RAFTS3 was compared with that of BLASTp version 2.2.26+, USEARCH and PAUDA. Tests were performed using Linux CentOS 6.5 on a Desktop AMD Six-Core 3.5Ghz processor with 8Gb of RAM, configuration details for each test are explained on each results section.

## RESULTS AND DISCUSSION

### Parameters selection analysis

To determine the default parameters to be used by RAFTS3 for the candidate selection step, sets of 1 to 20 k-mers with 4, 5, 6 or 7 amino acid residues were evaluated using the NR database. A subset of 1000 protein sequences randomly selected from NR was used as query. Two criteria were considered to define the RAFTS3 configuration settings: the running time to search 1000 queries (Figure 3.A); and the number of queries with second best hit with relative score higher than 0.3 (Figure 3.B). The best hit was disregarded since that always corresponds to the query sequence.

**Figure 2.**
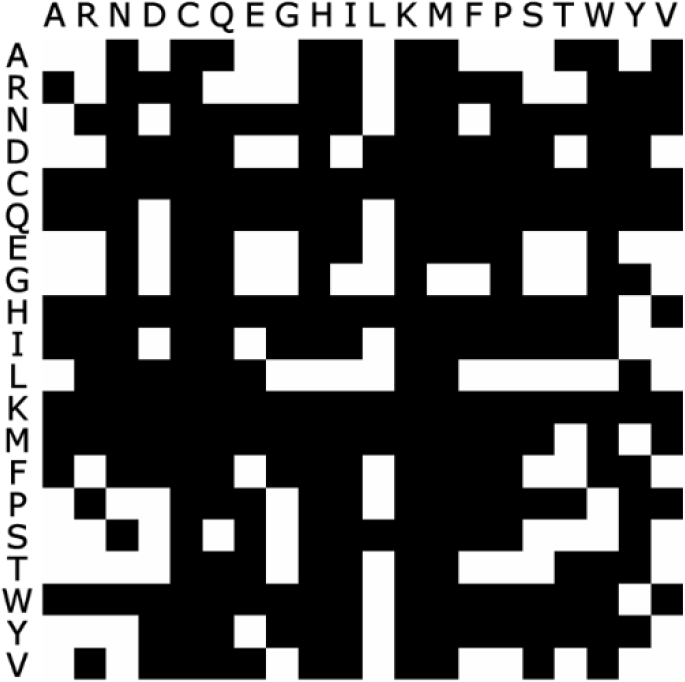
**Binary co-occurrence matrix (BCOM)**. Co-occurrence matrix of a protein sequence. White squares represent the occurrence of amino acid pairs; black squares represent non-occurrence.

**Figure. 3.**
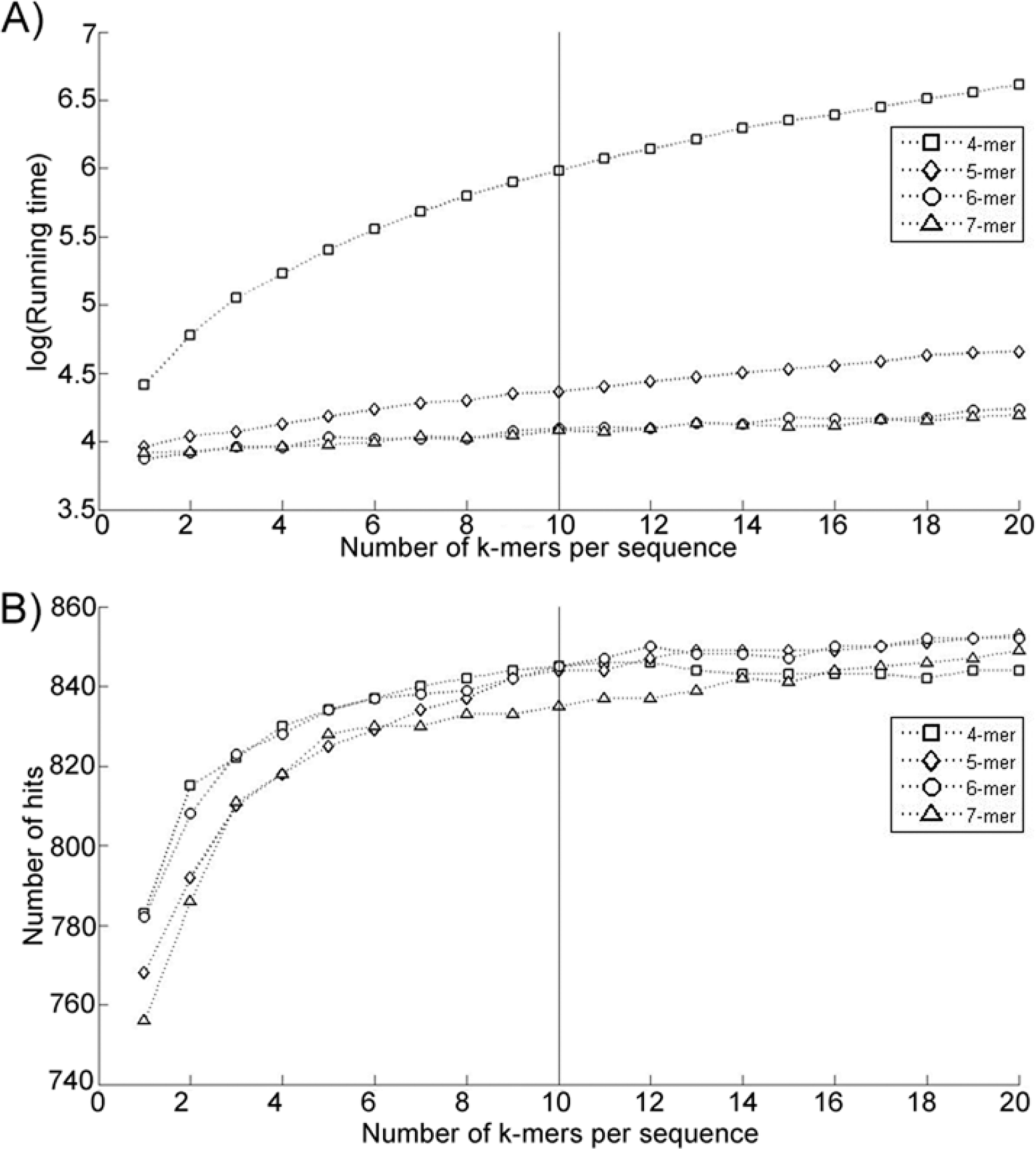
**Parameters selection and configuration testing**. Comparison of different k-mer sets. The comparison was made by analyzing the second best hit of the search results of 1000 sequences randomly selected. The number of k-mers ranged from 1 to 20 and their lengths were 4, 5, 6 and 7 amino acid residues. A) Shows the logarithm of the running time in seconds to search 1000 queries for each configuration. B) Shows the number of queries with second best hit with relative score over 0.3.

The purpose of this procedure was to find the number and size of k-mers to be adopted as default parameters to carry out searches with RAFTS3. This analysis showed that the running times were lower using k-mer sizes of 6 and 7 residues and the number of hits with relative score higher than 0.3 reached a plateau with sets of 10 k-mers per sequence. Thereby, the following parameters were chosen as default: 10 k-mers of 6 amino acid residues per sequence.

### Comparison of RAFTS3 with BLASTp

The sensitivity and running time of the RAFTS3 was compared with BLASTp using 1000 sequences randomly selected from a newer version of NR. These sequences were absent inthe database used for search tests and represent sequences from more than 650 different organisms. This comparison simulates an automated annotation task, thus BLASTp and RAFTS3 were configured only to report the best hit for comparison. The sensitivity was evaluated as the number of similar sequences retrieved and the processing time spent in the search by both tools. The number of sequences retrieved by BLASTp was considered as the gold standard, representing 100% of the results.

RAFTS3 showed results from 77% to 95% of sensitivity compared with BLAST p when searching UniRef50 database, from 86% to 95% when searching the Pfam database and from 89% to 97% when searching the NR database, depending on the threshold of the score (Table 2). RAFTS3 showed to be more than 300 times faster than BLASTp when searching in the larger database.

**Table 2.** Performance comparison of similarity search tools on the same query dataset (1000 sequences) against different protein databases

| Database | Total sequences | Total aa | Running time |  | Percentage of sequences over relative score threshold found by RAFTS3 compared with BLASTp |  |  |  |  |  |  |
| --- | --- | --- | --- | --- | --- | --- | --- | --- | --- | --- | --- |
|  |  |  | BLASTp | RAFTS3 | 0.3 | 0.4 | 0.5 | 0.6 | 0.7 | 0.8 | 0.9 |
| UniRef50 | 6,784,251 | 2,189,361,886 | 120m16.516s | 0m47.209s | 81% | 80% | 77% | 80% | 87% | 92% | 95% |
| Pfam | 15,929,002 | 5,169,768,107 | 262m22.350s | 0m52.832s | 86% | 91% | 92% | 93% | 95% | 95% | 94% |
| NR | 19,689,576 | 6,752,058,980 | 362m1.048s | 1m7.514s | 89% | 92% | 93% | 95% | 96% | 97% | 96% |

The total number of sequences and amino acids included in each database are shown in the “Total sequences” and “Total aa” columns, respectively. The ratio RAFTS3/BLASTp gives the fraction of RAFTS3 hits over BLASTp hits for theindicated relative score thresholds.

To illustrate the differences between the RAFTS3 and BLASTp hits, three different proteins were searched against the NR database: the pyrR (UniProtAC P39765) of Bacillus subtilis that regulates the transcription of the pyrimidine nucleotide (pyr) operon; the PRNP (UniProtAC P04165) of Homo sapiens related with neuronal development and synaptic plasticity; and the PSG1 (UniProtAC P11464) of Homo sapiens related with female pregnancy. The top 10 hits found by each were selected and the E-value and the relative scores were calculated for comparison. The results showed that, despite some differences, RAFTS3 performed similarly to BLASTp (Table3, 4 and 5).

**Table 3.** Comparison of top 10 of BLASTp and RAFTS3 hits of Bacillus subtilis PyrR protein

| RAFTS3 |  |  | BLASTp |  |  |
| --- | --- | --- | --- | --- | --- |
| Query - UniProtAC: P39765 Bifunctional protein PyrR OS=*Bacillus subtilis* |  |  |  |  |  |
| Relative Score | E-Value | Subject Sequence | Relative Score | E-value | Subject Sequence |
| 1 | 1.6941e-50 | gi\|16078611\|ref\|NP_389430.1\|<br>bifunctional pyrimidine regulatory<br>protein PyrR/uracil<br>phosphoribosyltransferase<br>[*Bacillus subtilis* subsp. *subtilis* str.<br>168] | 1 | 1.6941e-50 | gi\|16078611\|ref\|NP_389430.1\|<br>bifunctional pyrimidine regulatory<br>protein PyrR/uracil<br>phosphoribosyltransferase<br>[*Bacillus subtilis* subsp. *subtilis* str.<br>168] |
| 0.991274 | 4.9554e-50 | gi\|296331123\|ref\|ZP_06873597.1\|<br>bifunctional pyrimidine regulatory<br>protein PyrR uracil<br>phosphoribosyltransferase<br>[*Bacillus subtilis* subsp. *spizizenii*<br>ATCC 6633] | 0.991274 | 4.9554e-50 | gi\|296331123\|ref\|ZP_06873597.1\|<br>bifunctional pyrimidine regulatory<br>protein PyrR uracil<br>phosphoribosyltransferase<br>[*Bacillus subtilis* subsp. *spizizenii*<br>ATCC 6633] |
| 0.958988 | 2.5129e-48 | gi\|1373160\|gb\|AAB57770.1\|<br>PyrR, partial<br>[*Bacillus subtilis* subsp. *subtilis* str.<br>168] | 0.973822 | 4.24e-49 | gi\|398304125\|ref\|ZP_10507711.1\|<br>bifunctional pyrimidine regulatory<br>protein PyrR uracil<br>phosphoribosyltransferase<br>[*Bacillus vallismortis* DV1-F-3] |
| 0.973822 | 4.24e-49 | gi\|398304125\|ref\|ZP_10507711.1\|<br>bifunctional pyrimidine regulatory<br>protein PyrR uracil<br>phosphoribosyltransferase<br>[*Bacillus vallismortis* DV1-F-3] | 0.954625 | 4.4966e-48 | gi\|154685963\|ref\|YP_001421124.1\|<br>bifunctional pyrimidine regulatory<br>protein PyrR uracil<br>phosphoribosyltransferase<br>[*Bacillus amyloliquefaciens* FZB42] |
| 0.954625 | 4.4966e-48 | gi\|154685963\|ref\|YP_001421124.1\|<br>bifunctional pyrimidine regulatory<br>protein PyrR uracil<br>phosphoribosyltransferase<br>[*Bacillus amyloliquefaciens* FZB42] | 0.951134 | 6.9078e-48 | gi\|311068068\|ref\|YP_003972991.1\|<br>bifunctional pyrimidine regulatory<br>protein PyrR/uracil<br>phosphoribosyltransferase<br>[*Bacillus atrophaeus* 1942] |
| 0.953752 | 5.006e-48 | gi\|375362191\|ref\|YP_005130230.1\|<br>bifunctional pyrimidine regulatory<br>protein PyrR/uracil<br>phosphoribosyltransferase<br>[*Bacillus amyloliquefaciens* subsp.<br>*plantarum* CAU B946] | 0.958988 | 2.5129e-48 | gi\|1373160\|gb\|AAB57770.1\|<br>PyrR, partial<br>[*Bacillus subtilis* subsp. *subtilis* str.<br>168] |
| 0.951134 | 6.9078e-48 | gi\|311068068\|ref\|YP_003972991.1\|<br>bifunctional pyrimidine regulatory<br>protein PyrR/uracil<br>phosphoribosyltransferase<br>[*Bacillus atrophaeus* 1942] | 0.953752 | 5.006e-48 | gi\|375362191\|ref\|YP_005130230.1\|<br>bifunctional pyrimidine regulatory<br>protein PyrR/uracil<br>phosphoribosyltransferase<br>[*Bacillus amyloliquefaciens* subsp.<br>*plantarum* CAU B946] |
| 0.904887 | 2.0525e-45 | gi\|52080149\|ref\|YP_078940.1\|<br>bifunctional pyrimidine regulatory<br>protein PyrR uracil<br>phosphoribosyltransferase<br>[*Bacillus licheniformis* DSM 13 =<br>ATCC 14580] | 0.904887 | 2.0525e-45 | gi\|52080149\|ref\|YP_078940.1\|<br>bifunctional pyrimidine regulatory<br>protein PyrR uracil<br>phosphoribosyltransferase<br>[*Bacillus licheniformis* DSM 13 =<br>ATCC 14580] |
| 0.877836 | 5.6563e-44 | gi\|389573331\|ref\|ZP_10163406.1\|<br>bifunctional protein PyrR<br>[*Bacillus aerophilus* KACC 16563] | 0.877836 | 5.6563e-44 | gi\|389573331\|ref\|ZP_10163406.1\|<br>bifunctional protein PyrR<br>[*Bacillus aerophilus* KACC 16563] |
| 0.873473 | 9.6739e-44 | gi\|157692227\|ref\|YP_001486689.1\|<br>pyrR gene product<br>[*Bacillus pumilus* SAFR-032] | 0.872600 | 1.077e-43 | gi\|194014677\|ref\|ZP_03053294.1\|<br>bifunctional protein PyrR<br>[*Bacillus pumilus* ATCC 7061] |

Comparison of the ten first results of RAFTS3 and BLASTp searching the Bacillus subtilis PyrR protein against the NR database. The subject sequences are ordered by each software default criteria.

**Table 4.**
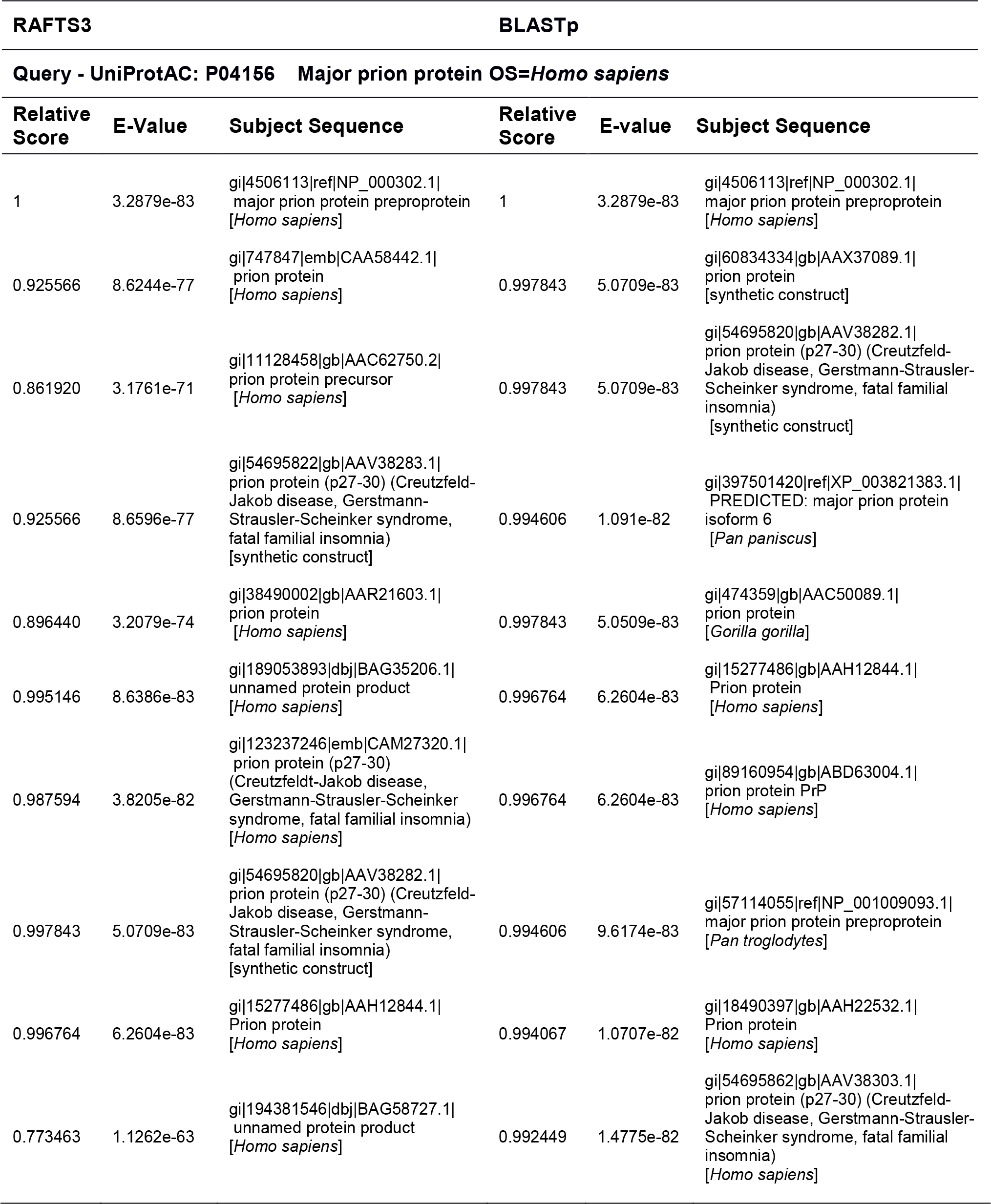
Comparison of top 10 of BLASTp and RAFTS3 hits of Homo sapiens Prion protein

| RAFTS3 |  |  | BLASTp |  |  |
| --- | --- | --- | --- | --- | --- |
| Query - UniProtAC: P04156 Major prion protein OS=*Homo sapiens* |  |  |  |  |  |
| Relative Score | E-Value | Subject Sequence | Relative Score | E-value | Subject Sequence |
| 1 | 3.2879e-83 | gi\|4506113\|ref\|NP_000302.1\| major prion protein preproprotein [*Homo sapiens*] | 1 | 3.2879e-83 | gi\|4506113\|ref\|NP_000302.1\| major prion protein preproprotein [*Homo sapiens*] |
| 0.925566 | 8.6244e-77 | gi\|747847\|emb\|CAA58442.1\| prion protein [*Homo sapiens*] | 0.997843 | 5.0709e-83 | gi\|60834334\|gb\|AAX37089.1\| prion protein [synthetic construct] |
| 0.861920 | 3.1761e-71 | gi\|11128458\|gb\|AAC62750.2\| prion protein precursor [*Homo sapiens*] | 0.997843 | 5.0709e-83 | gi\|54695820\|gb\|AAV38282.1\| prion protein (p27-30) (Creutzfeld-Jakob disease, Gerstmann-Strausler-Scheinker syndrome, fatal familial insomnia) [synthetic construct] |
| 0.925566 | 8.6596e-77 | gi\|54695822\|gb\|AAV38283.1\| prion protein (p27-30) (Creutzfeld-Jakob disease, Gerstmann-Strausler-Scheinker syndrome, fatal familial insomnia) [synthetic construct] | 0.994606 | 1.091e-82 | gi\|397501420\|ref\|XP_003821383.1\| PREDICTED: major prion protein isoform 6 [*Pan paniscus*] |
| 0.896440 | 3.2079e-74 | gi\|38490002\|gb\|AAR21603.1\| prion protein [*Homo sapiens*] | 0.997843 | 5.0509e-83 | gi\|474359\|gb\|AAC50089.1\| prion protein [*Gorilla gorilla*] |
| 0.995146 | 8.6386e-83 | gi\|189053893\|dbj\|BAG35206.1\| unnamed protein product [*Homo sapiens*] | 0.996764 | 6.2604e-83 | gi\|15277486\|gb\|AAH12844.1\| Prion protein [*Homo sapiens*] |
| 0.987594 | 3.8205e-82 | gi\|123237246\|emb\|CAM27320.1\| prion protein (p27-30) (Creutzfeldt-Jakob disease, Gerstmann-Strausler-Scheinker syndrome, fatal familial insomnia) [*Homo sapiens*] | 0.996764 | 6.2604e-83 | gi\|89160954\|gb\|ABD63004.1\| prion protein PrP [*Homo sapiens*] |
| 0.997843 | 5.0709e-83 | gi\|54695820\|gb\|AAV38282.1\| prion protein (p27-30) (Creutzfeld-Jakob disease, Gerstmann-Strausler-Scheinker syndrome, fatal familial insomnia) [synthetic construct] | 0.994606 | 9.6174e-83 | gi\|57114055\|ref\|NP_001009093.1\| major prion protein preproprotein [*Pan troglodytes*] |
| 0.996764 | 6.2604e-83 | gi\|15277486\|gb\|AAH12844.1\| Prion protein [*Homo sapiens*] | 0.994067 | 1.0707e-82 | gi\|18490397\|gb\|AAH22532.1\| Prion protein [*Homo sapiens*] |
| 0.773463 | 1.1262e-63 | gi\|194381546\|dbj\|BAG58727.1\| unnamed protein product [*Homo sapiens*] | 0.992449 | 1.4775e-82 | gi\|54695862\|gb\|AAV38303.1\| prion protein (p27-30) (Creutzfeld-Jakob disease, Gerstmann-Strausler-Scheinker syndrome, fatal familial insomnia) [*Homo sapiens*] |

Comparison of the ten first results of RAFTS3 and BLASTp searching the Homo sapiensPrion protein against the NR database. The subject sequences are ordered by each software default criteria.

**Table 5.** Comparison of top 10 of BLASTp and RAFTS3 hits of Homo sapiens PSG1 protein

| RAFTS3 |  |  | BLASTp |  |  |
| --- | --- | --- | --- | --- | --- |
| Query - UniProtAC: P11464 Pregnancy-specific beta-1-glycoprotein 1 OS=*Homo sapiens* |  |  |  |  |  |
| Relative Score | E-Value | Subject Sequence | Relative Score | E-value | Subject Sequence |
| 1 | 3.9757e-128 | gi\|296317345\|ref\|NP_001171754.1\| pregnancy-specific beta-1-glycoprotein 1 isoform 2 precursor [*Homo sapiens*] | 1 | 3.9757e-128 | gi\|296317345\|ref\|NP_001171754.1\| pregnancy-specific beta-1-glycoprotein 1 isoform 2 precursor [*Homo sapiens*] |
| 0.996463 | 1.1629e-127 | gi\|190645\|gb\|AAA36515.1\| pregnancy-specific glycoprotein-1a [*Homo sapiens*] | 0.996463 | 1.1629e-127 | gi\|190645\|gb\|AAA36515.1\| pregnancy-specific glycoprotein-1a [*Homo sapiens*] |
| 0.985497 | 3.31e-126 | gi\|306797\|gb\|AAA52602.1\| pregnancy-specific beta-glycoprotein c [*Homo sapiens*] | 0.985497 | 3.2249e-126 | gi\|296317348\|ref\|NP_001171755.1\| pregnancy-specific beta-1-glycoprotein 1 isoform 3 precursor [*Homo sapiens*] |
| 0.985497 | 3.2249e-126 | gi\|296317348\|ref\|NP_001171755.1\| pregnancy-specific beta-1-glycoprotein 1 isoform 3 precursor [*Homo sapiens*] | 0.985497 | 3.31e-126 | gi\|306797\|gb\|AAA52602.1\| pregnancy-specific beta-glycoprotein c [*Homo sapiens*] |
| 0.981606 | 1.0502e-125 | gi\|306791\|gb\|AAA52590.1\| pregnancy-specific beta-1-glycoprotein [*Homo sapiens*] | 0.989034 | 1.1263e-126 | gi\|21361392\|ref\|NP_008836.2\| pregnancy-specific beta-1-glycoprotein 1 isoform 1 precursor [*Homo sapiens*] |
| 0.989034 | 1.1263e-126 | gi\|21361392\|ref\|NP_008836.2\| pregnancy-specific beta-1-glycoprotein 1 isoform 1 precursor [*Homo sapiens*] | 0.981606 | 1.0502e-125 | gi\|306791\|gb\|AAA52590.1\| pregnancy-specific beta-1-glycoprotein [*Homo sapiens*] |
| 0.985497 | 3.2945e-126 | gi\|190653\|gb\|AAA36517.1\| pregnancy-specific glycoprotein-1d [*Homo sapiens*] | 0.985497 | 3.2945e-126 | gi\|190653\|gb\|AAA36517.1\| pregnancy-specific glycoprotein-1d [*Homo sapiens*] |
| 0.985143 | 3.6678e-126 | gi\|190591\|gb\|AAA36511.1\| pregnancy-specific beta-1-glycoprotein [*Homo sapiens*] | 0.985143 | 3.6678e-126 | gi\|190591\|gb\|AAA36511.1\| pregnancy-specific beta-1-glycoprotein [*Homo sapiens*] |
| 0.569155 | 1.8476e-71 | gi\|3287447\|gb\|AAC25485.1\| PSGIIA-a [*Homo sapiens*] | 0.948709 | 2.2829e-121 | gi\|14250018\|gb\|AAH08405.1\| PSG4 protein [*Homo sapiens*] |
| 0.948709 | 2.2829e-121 | gi\|14250018\|gb\|AAH08405.1\| PSG4 protein [*Homo sapiens*] | 0.949416 | 1.8419e-121 | gi\|332855947\|ref\|XP_512709.3\| PREDICTED: pregnancy-specific beta-1-glycoprotein 1 isoform 3 [*Pan troglodytes*] |

Comparison of the ten first results of RAFTS3 and BLASTp searching theHomo sapiensPSG1 protein against the NR database. The subject sequences are ordered byeach software default criteria.

To compare the ranking order of sequences given by BLASTp and RAFTS3, 1000 sequences were randomly selected from the dataset to be used as query against the NR database.The position of RAFTS3 best hits were scored among BLASTp top 50 hits and vice-versa. The results showed that 72% of the RAFTS3 best hits occurred within the first 10 BLASTp top results (Supplementary material Table S1), suggesting that sequences retrieved by RAFTS3 are in most the same or very closely related to that retrieved by BLASTp. To better illustrate the ranking differences between BLASTp and RAFTS3 the top 50 hits identified by RAFTS3 and BLASTp using 5 different proteins randomly selected from the test set as query to search against the NR database are shown in supplementary material Table S2. In all cases, BLASTp best hit was among the 10 best hits of RAFTS3. Interestingly, for steroidogenic factor 1 isoform X2 RAFTS3 top hit had a higher relative score that of BLASTp (Table S2).

### Comparison of RAFTS3 with USEARCH

USEARCH provides freely only a version with limited use of resources, the complete version of requires a paid license. Thereby, we chose to use the small COG [26] database to compare RAFTS3 and USEARCH performances. In this test, USEARCH was faster and more accurate than RAFTS3. However, due to the limitations of the free version, it was not possible to evaluate USEARCH performance searching large databases. It is possible to anticipate that memory consumption of USEARCH will be more than 40Gb for the NR database, while RAFTS3 uses 20 times less. Also RAFTS3 runtime is not much affected by the database size and the sensitivity tends to increase. These considerations indicate that the use of RAFTS3 may be advantageous over USEARCH when searching large databases.

### Comparison of RAFTS3 with PAUDA

To compare RAFTS3 with PAUDA an executable was developed to translate DNA sequences in all 6 frames to search on a protein database. We called it RAFST3x (in analogy to BLASTx). The tests were performed comparing 1000 sequences randomly selected from the NT database (lengths from 50 to 3000 pb) against the UniRef50 database. RAFTS3x was 7%faster than PAUDA and more sensitive, yielding twice as many hits above the threshold relative score (Table 6).

**Table 6.**
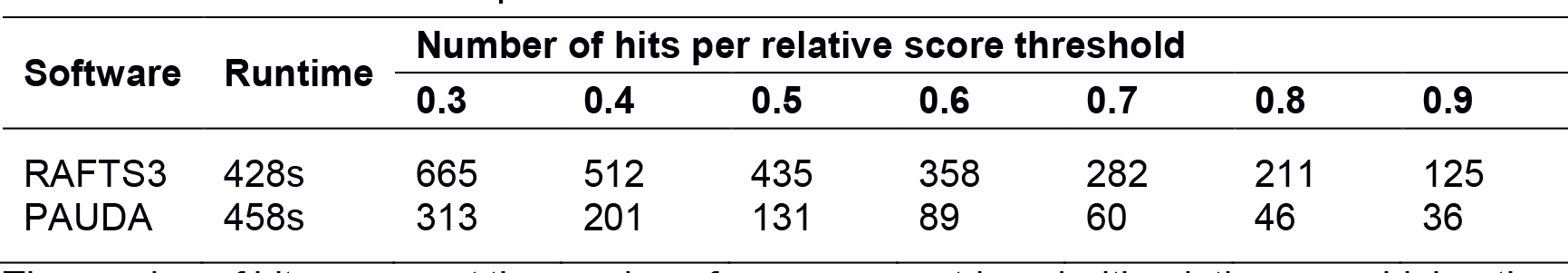
Performance comparison of RAFTS3 and PAUDA

The number of hits represent the number of sequences retrieved with relative score higher than the relative score threshold.

### Alignment information

The performance advantage of RAFTS3 relies on the comparison of sequences without the need of alignment. The measure used is based on the BCOM’s comparison that have some relationship with an alignment score. It’s possible to perform local alignments on the hits reported by RAFTS3 using the Smith-Waterman algorithm, however this adds an additional cost on time. The runtime for RAFTS3 configurations using from 0 to 100 alignments to search the 1000 sequences against the NR database varied from 40 seconds to 17 minutes. Thus this option must be used wisely.

## CONCLUSIONS

RAFTS3 uses an aggressive filter approach with a fast comparison method based on BCOMs. Due to the limitation of the free version of USEARCH, comparisons for searches against large databases could not be performed. The comparison of RAFTS3 with BLASTp showed that RAFTS3 could be used to achieve fast protein similarity searches with a small loss of sensitivity. The sensitivity compared to BLAST p increases with the sequence similarity. RAFTS3 also shows a minimal loss on performance when challenged with larger databases in comparison with BLASTp, as judged by the increase in time to search on UniRef50 compared to NR (almost 3 times as large), the running time for RAFTS3 increase twice while BLAST p increased thrice. Thus RAFTS3 could be especially advantageous when using large databases with many sequences being queried. As the database increases, the filtering options can be made more stringent avoiding the increase of the number of candidate sequences selected and, consequently, of memory usage.

We have demonstrated that the RAFTS3 can perform high-speed protein search comparisons locally using a desktop computer or laptop. RAFTS3 is being used in tasks as genome annotation by our Bioinformatics group at the Federal University of Parana with success and presents a good solution for protein sequence data mining.

## DECLARATIONS

### Availability and requirements

Project name: RAFTS3
Project home page: https://sourceforge.net/projects/rafts3/
Operating system(s): Linux, Windows Programming language: MATLAB (R2012a)
Other requirements: MCR MATLAB Compiler Runtime v7.17 (only for compiled version) License: Source code and binaries freely available under the BSD License Any restrictions to use by non-academics: None

## Authors’ contributions

RAV implemented the software, validated the results and wrote the manuscript. RTR designed the study and developed the prototype. FOP, VAW, JNM and EMS contributed to the concepts and revised the manuscript. DG contributed to the concepts. JHT contributed to the testing. All authors read and approved the final manuscript.

## SUPPORTING INFORMATION

**Table S1.**
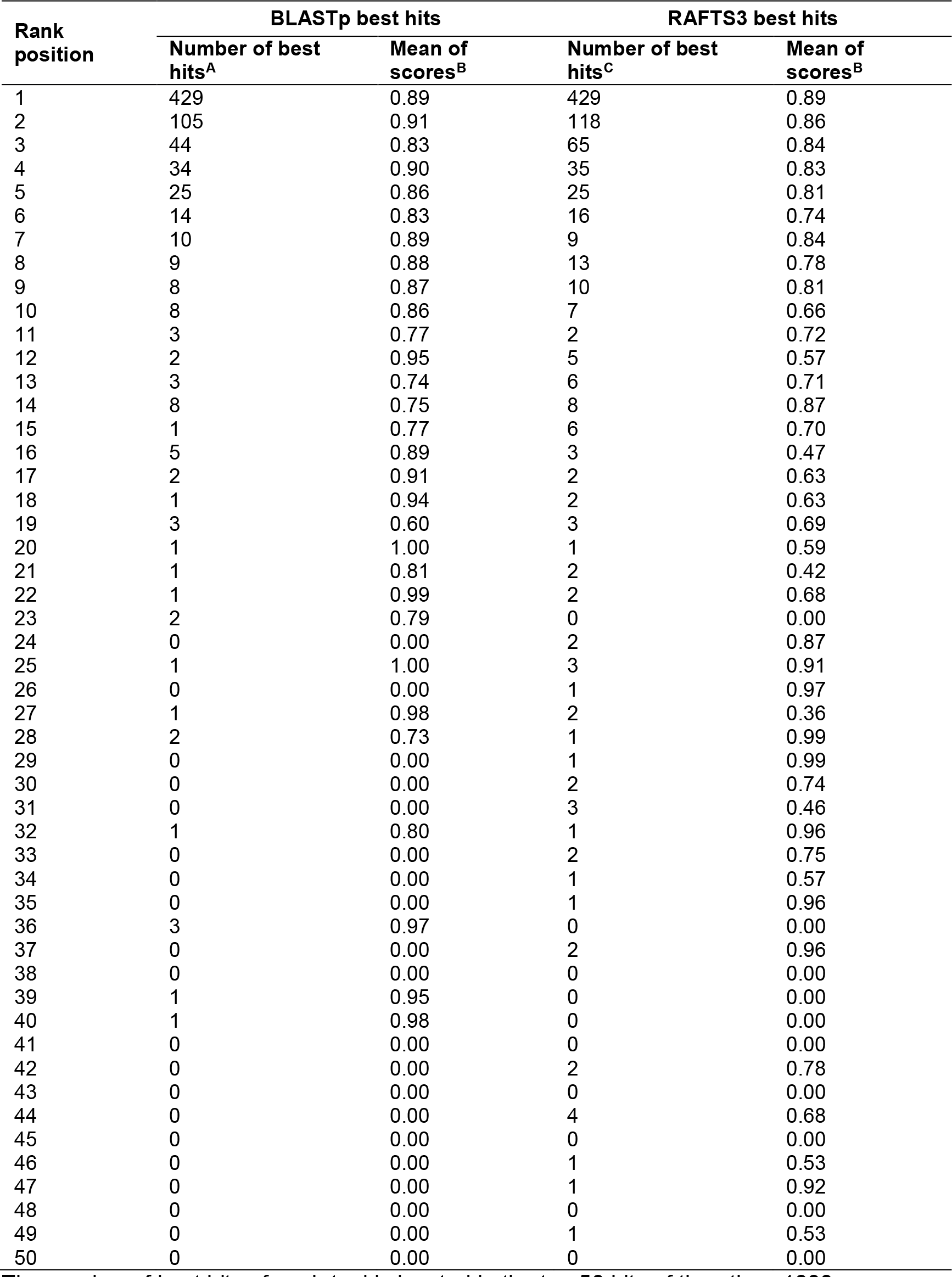
Comparison of BLASTp and RAFTS3 hits ranking.

The number of best hits of each tool is located in the top 50 hits of the other. 1000 sequences of the data set were used as queries for searching NR database with BLASTp or RAFTS3, and the number of best hit of each tool was scored for each rank position of the other tool. For example, 429 best hits of BLASTp are also best hits of RAFTS3, while 105 best hits of RAFTS3 are second best hits of BLASTp.
A Number of RAFTS3 best hits occurring in the indicated BLASTp rank position.
B Average relative score of the best hits occurring in the indicated rank position.
C Number of BLASTp best hits occurring in the indicated RAFTS3 rank position.

**Table S2.**
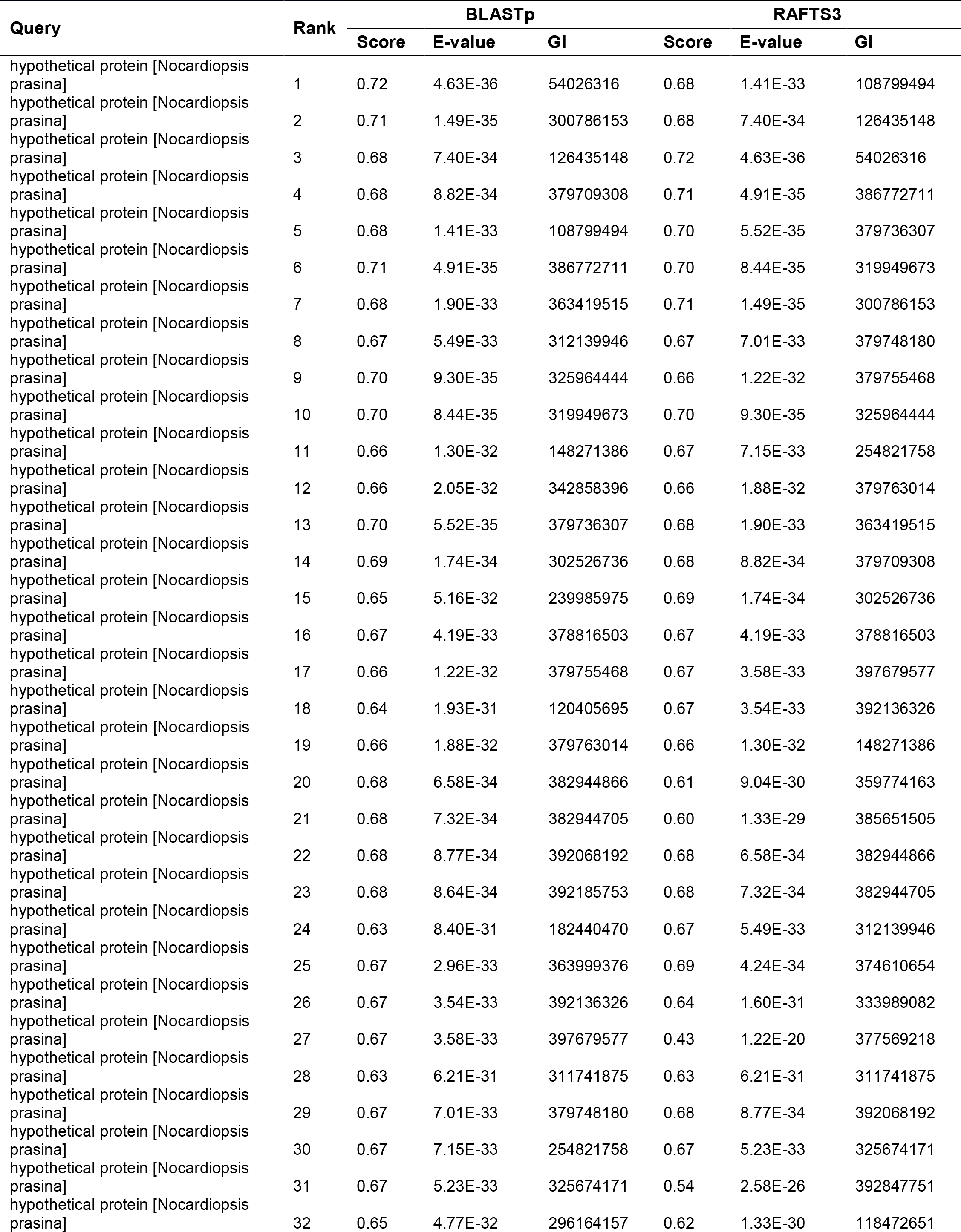
Problem-solving strategies that separate qualitative/pictorial steps from mathematical steps.

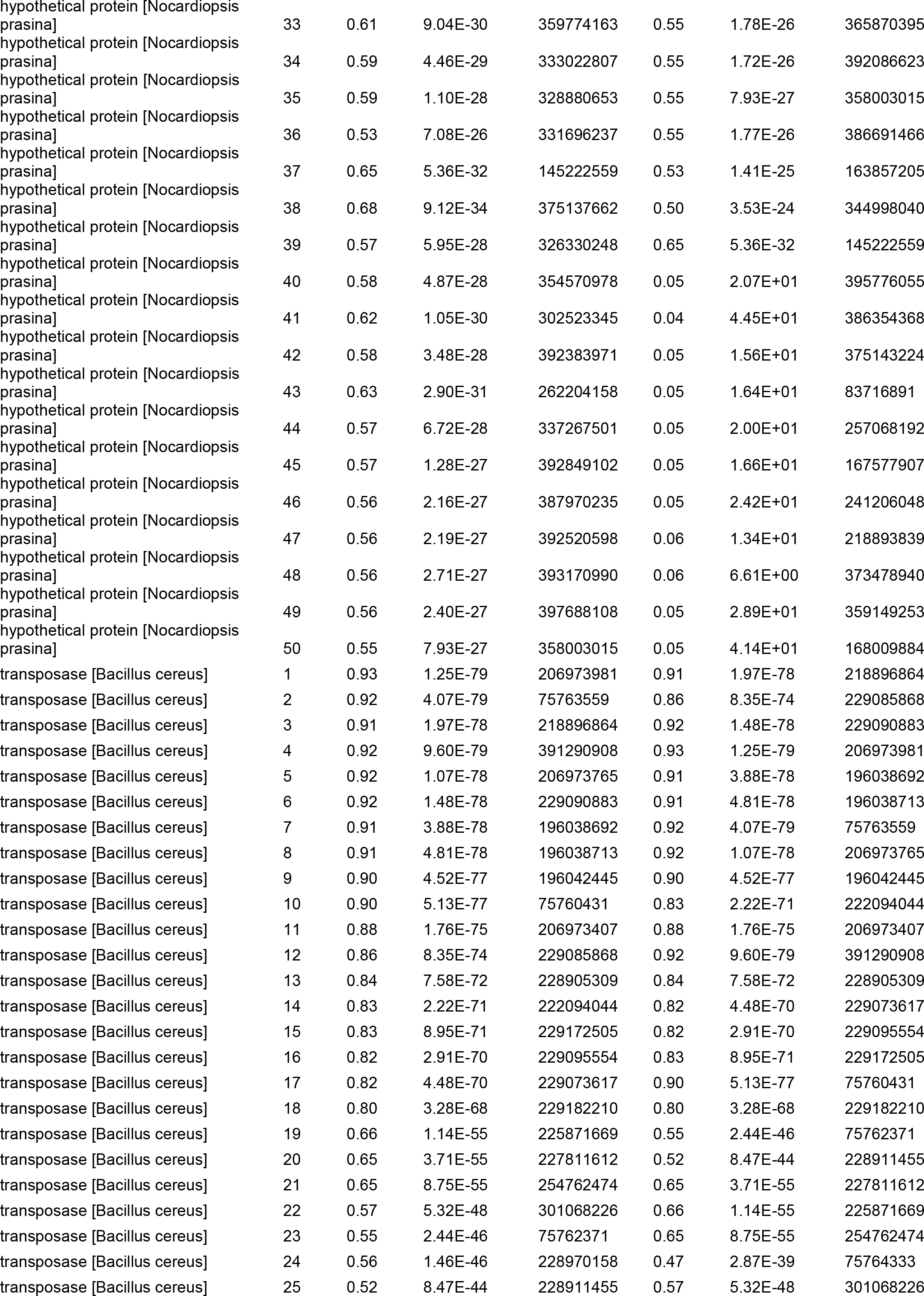

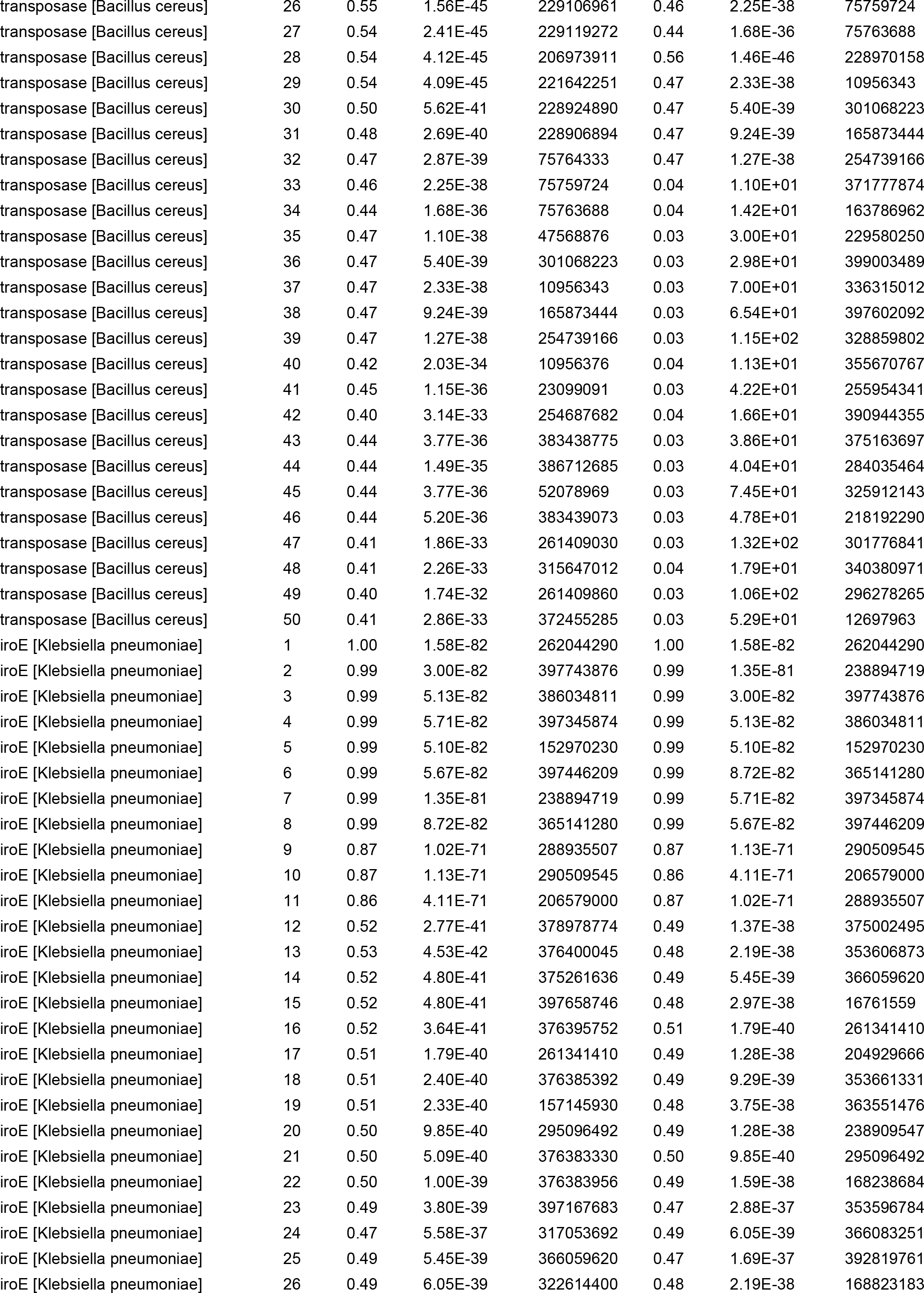

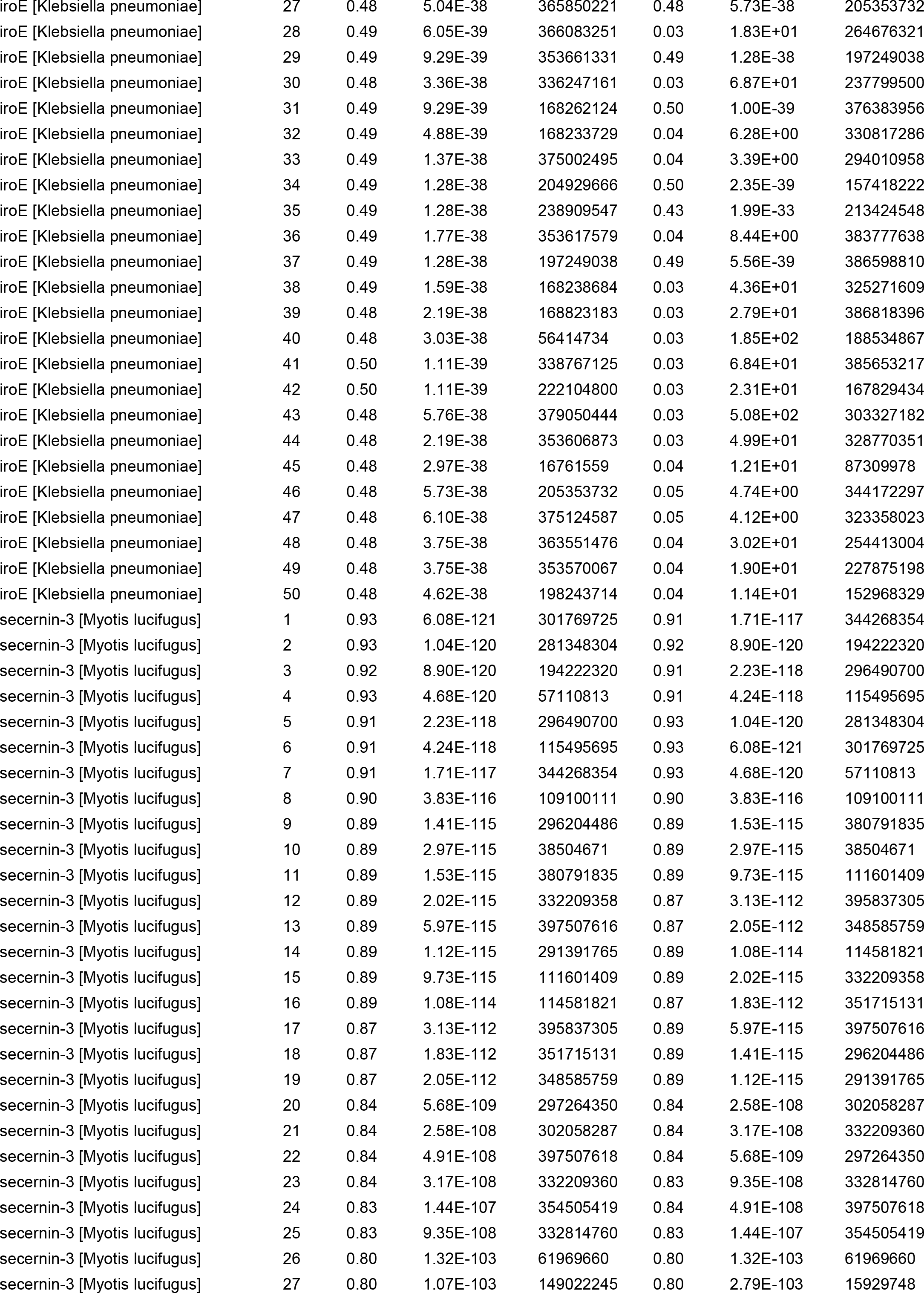

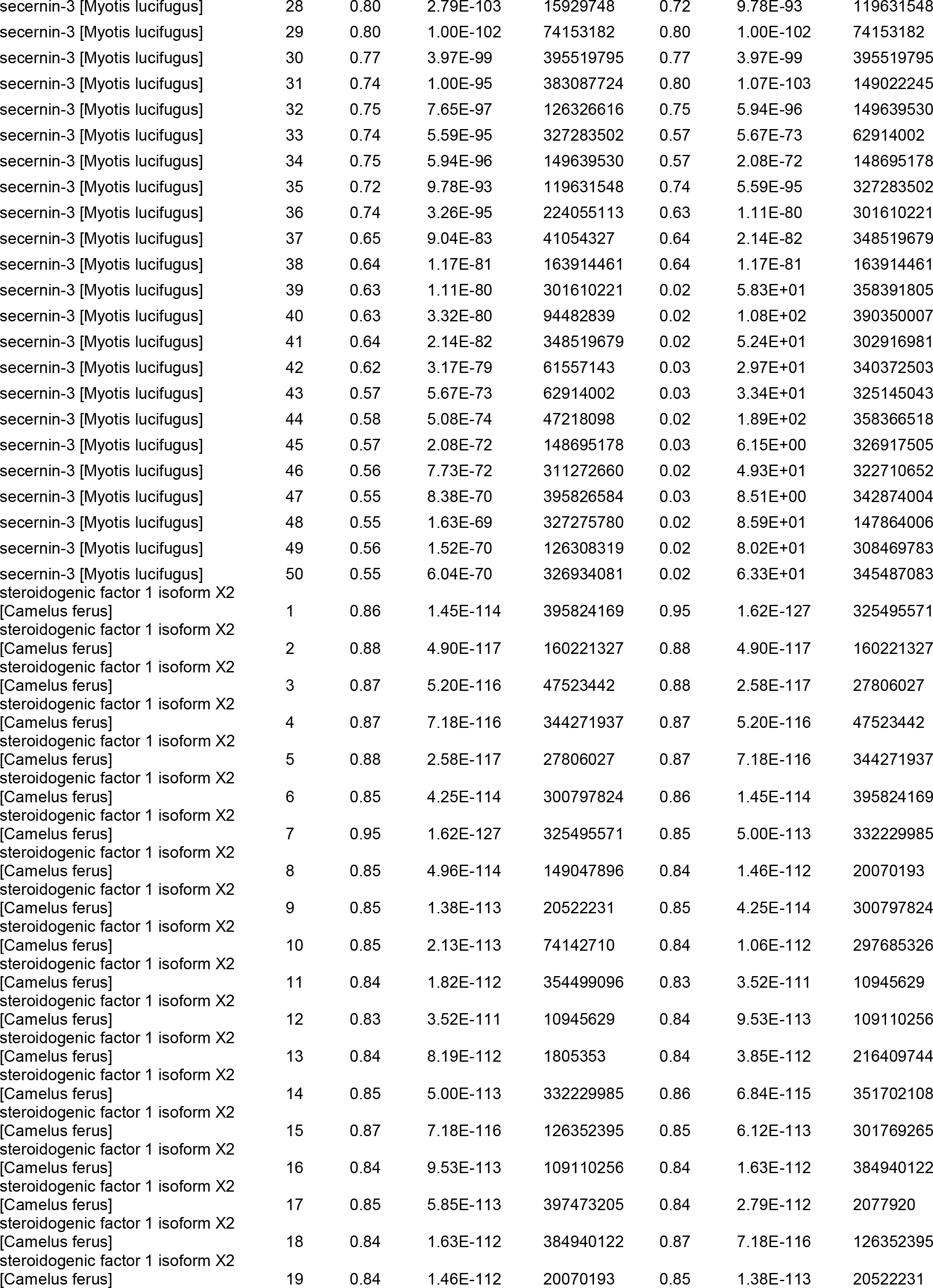

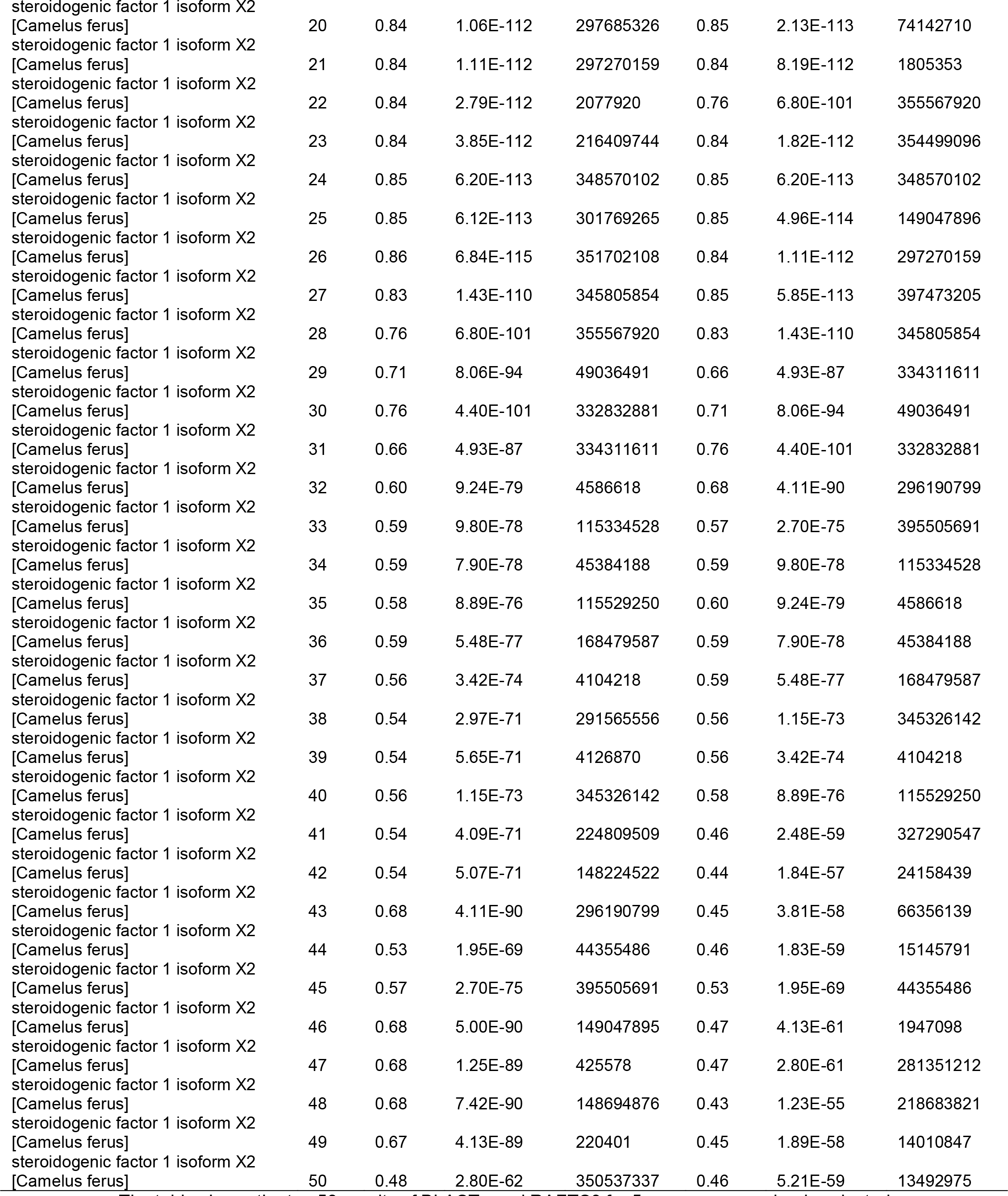

The table shows the top 50 results of BLASTp and RAFTS3 for 5 sequences randomly selected from the test dataset compared against NR. The subject sequences are indicated by their GI number and ordered by the default criteria of each tool; the relative score of each one was calculated.

